# Comparative analysis of whole genome amplification kits for single-cell genome analysis

**DOI:** 10.1101/2025.10.30.685509

**Authors:** Abrahan Hernández-Hernández, Pontus Höjer, Michelle Ljungmark, Anastasios Glaros, Eunkyoung Choi, Julia Hauenstein, Nicolai Frengen, Susanne Björnerfeldt, Camilla Enström, Robert Månsson, Jessica Nordlund, Anja Mezger

## Abstract

Whole genome amplification (WGA) enables detailed analysis of genomic heterogeneity at the cellular level. Here, we systematically compare two commercially available WGA technologies on single cells; Multiple Displacement Amplification in the Qiagen REPLI-g kit (MDA/REPLI-g) and Primary Template-directed Amplification in the BioSkryb ResolveDNA kit (PTA/ResolveDNA). Beyond quality control and genomic profiling, we assessed a cost-effective array-based approach as a complement to low-pass sequencing. Consistent with available benchmarks, PTA/ResolveDNA sequencing libraries consistently outperformed MDA/REPLI-g across quality metrics, including genome coverage breadth and uniformity. CNV calls from PTA-cells showed higher accuracy and sensitivity, including reliable detection of small CNVs at low sequencing depth. SNP array-based analysis corroborated sequencing-based findings, and additionally provided allelic information. Notably, CNV detection from SNP arrays matched the performance of low-pass sequencing, supporting its utility as an cost-effective approach for WGA quality control and single-cell CNV profiling.

## INTRODUCTION

Single-cell whole genome sequencing (scWGS) has revolutionized our understanding of cellular heterogeneity, enabling detailed insights into genetic variations at the single cell level (Zong et al. 2012; McConnell et al. 2013; Wang et al. 2014). One of the main challenges with scWGS is the limited amount of DNA material derived from a single cell (∼6 pg), necessitating whole genome amplification (WGA) for comprehensive genome analysis (Gawad et al. 2016).

An ideal WGA method for a single cell would faithfully and uniformly amplify the entire genome while preserving the original nucleotide sequence and allelic composition. In practice, however, the small amount of DNA present in a single cell requires amplification strategies that inevitably introduce technical biases, including uneven genome coverage, sequence errors, and amplification artifacts. To address these challenges, numerous WGA methods have been developed, each balancing amplification fidelity, and coverage uniformity differently. Among these, Multiple Displacement Amplification (MDA) has become one of the most widely adopted approaches (Dean et al. 2001; Zhang et al. 2001). The method uses random priming with a highly-fidelity, strand displacing phi29 polymerase to exponentially amplify DNA in a branching fashion. While MDA covers most of the genome and has a low error rate, it suffers from uneven coverage and amplification bias (Khoshkhoo et al. 2021; Estévez-Gómez et al. 2025). Other non-MDA methods such as PicoPlex, MALBAC, Ampli-1 and LIANTI employ amplification strategies that result in an improved genome coverage uniformity but with trade-offs in nucleotide-level fidelity (Khoshkhoo et al. 2021; Estévez-Gómez et al. 2025).

Primary Template-directed Amplification (PTA) was recently introduced as an evolution of the MDA principle, aiming to address several of the MDA-associated limitations. (Gonzalez-Pena et al. 2021). Similar to MDA, PTA utilizes random priming and Phi29 polymerase but incorporates exonuclease-resistant terminator nucleotides that restrict re-amplification of newly synthesized products, producing shorter amplicons while reducing amplification bias and enhancing coverage uniformity (Gonzalez-Pena et al. 2021; Middelkamp et al. 2023). Benchmarking studies comparing PicoPLEX, MDA/droplet MDA and PTA (Gonzalez-Pena et al. 2021; Jamal et al. 2021; Kalef-Ezra et al. 2024) have demonstrated improved genome coverage and variant detection performance for PTA, further establishing it as a promising approach for comprehensive single-cell genome analysis.

However, substantial differences may exist between commercial implementations of these technologies, and their performance in routine experimental workflows remains incompletely characterized. In particular, direct comparisons between widely used commercial kits, such as the MDA-based Qiagen REPLI-g Single Cell Kit (MDA/REPLI-g) and the PTA-based BioSkryb ResolveDNA Kit (PTA/ResolveDNA), remain limited. Here, we systematically compared amplification of single cells using these two kits across metrics including genome coverage breadth, amplification uniformity, amplification bias, and CNV detection accuracy. Furthermore, while previous studies have primarily focused on sequencing-based PTA performance metrics (Gonzalez-Pena et al. 2021; Middelkamp et al. 2023; Kalef-Ezra et al. 2024), we investigate genome-wide genotyping arrays as a complementary and cost-effective alternative to low-pass sequencing. By highlighting the strengths and weaknesses of each kit and analysis method, we aim to provide researchers with a comprehensive understanding of which technique is best suited for their specific scWGS applications.

## METHODS

### Cell culture and FACS sorting

Cultured MM.1S cells (a multiple myeloma cell-line) were incubated with propidium iodide (PI) and FACS sorted using the BD FACSAria IIu. Single or multiple live (PI-) cells were deposited using single-cell mode into 96-well plates containing the recommended buffer for the MDA/Repli-G and PTA/ResolveDNA kits (4 μL of scPBS and 3 μL of Cell Buffer, respectively). Cell sorting was performed on the same occasion and from the same sample of MM.1S cells to ensure minimal sample variation. For each method, 26 wells with single cells and two wells with 10 cells (bulk) were FACS sorted into the 96-well plate. In addition, two positive control wells with 1 ng of genomic DNA (provided with the BioSkryb ResolveDNA WGA Kit) in buffer and two empty wells (negative control containing only buffer) were included in each plate (Figure 1A).

**Figure 1.**
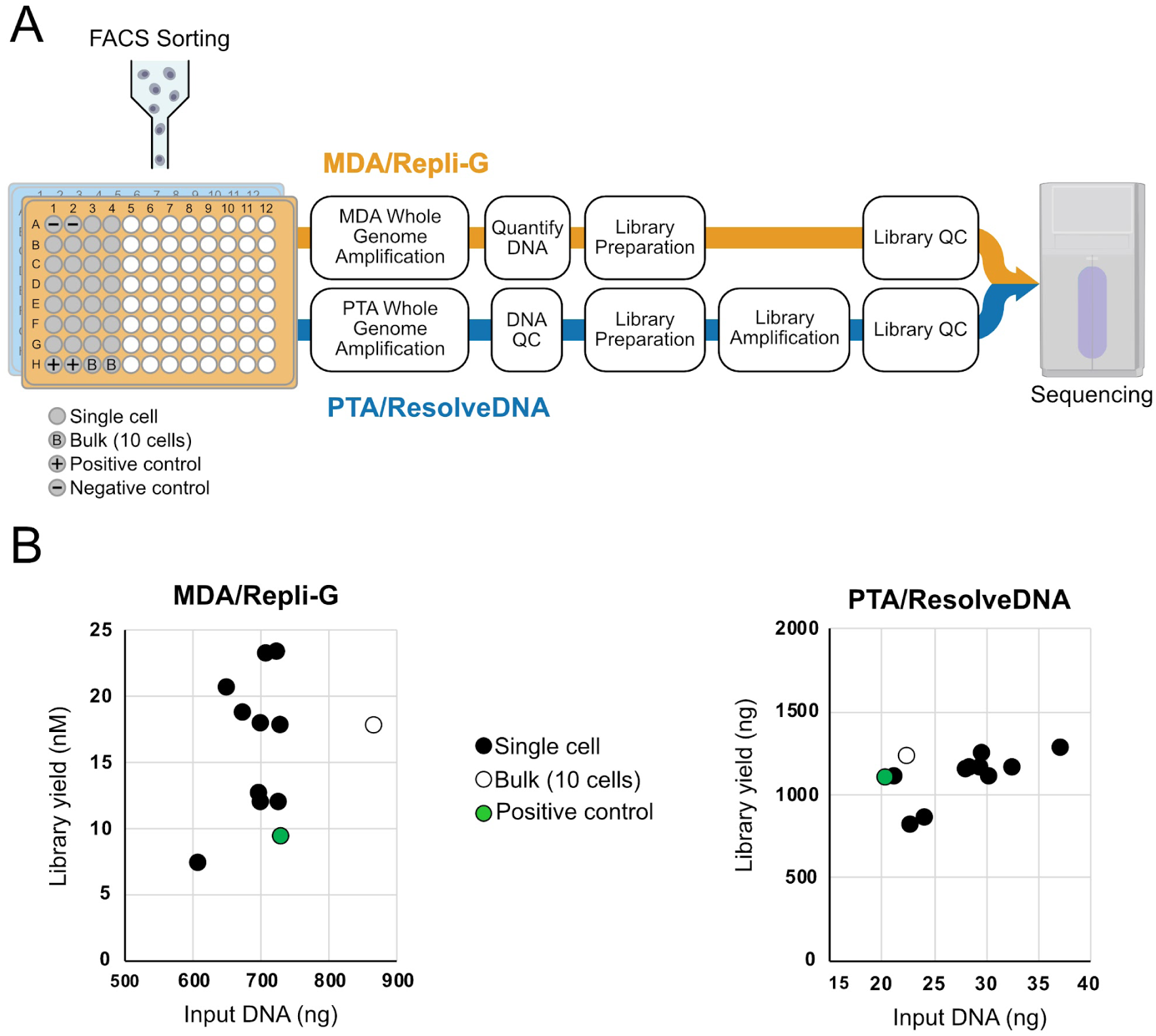
Plate layout, workflow and library yields for both of the tested scWGA kits. (**A**). Overview of sample processing with both MDA/REPLI-g and PTA/ResolveDNA kits. MM.1S cells were sorted into two fully skirted 96-well plates, each plate containing 26 single-cell wells and two bulk wells with ten cells. Two negative (buffer only), and two positive (1 ng extracted DNA) control wells were further included on both plates. Whole genome amplification was performed using the respective kit for each plate and ten random single cells plus one bulk from each underwent library preparation and sequencing. (**B**) Sequencing library yields (Y-axis) produced by the MDA/REPLI-g kit (left graph) and the PTA/ResolveDNA kit (right graph), using the WGA amplified by MDA and PTA, respectively. The amount of input DNA used to prepare sequencing libraries is displayed in the X axis.

### DNA amplification and library generation

WGA by MDA followed by library preparation was carried out with the QIAseq Single Cell Library kit with UDI, which combines the REPLI-g Advanced MDA for amplification, followed by enzymatic fragmentation and library ligation (Qiagen; Cat. nos. 1124559, 180318, and 1124693), whereas WGA by PTA with subsequent library preparation was performed using the ResolveDNA kit (BioSkryb; PN 100136, 100691, and 100940), following the manufacturer’s protocols. Following library preparation, ten randomly selected single-cell wells and one bulk well from each WGA method were subjected to qPCR quantification prior to sequencing.

### Illumina sequencing

The 22 quantified libraries were pooled and loaded into one full lane of a 25B flowcell (Illumina) at a final concentration of 190 pM with 1% PhiX. Sequencing was performed in a NovaSeq X Plus sequencer (Illumina) with the read configuration 151nt(R1)-10nt(I1)-10nt(I2)-151nt(R2).

### Genotyping

WGA amplified gDNA (200 ng) from the bulk and single-cell samples was genotyped using the Illumina Infinium Global Screening Array v3.0 (Illumina). Arrays were processed on an Illumina iScan system according to the manufacturer’s instructions. Initial data processing and genotype calling were performed in GenomeStudio (Illumina, v2.0.3), using Human Reference Genome build 38 (GRCh38).

### Illumina sequencing data analysis

Sequencing BCL files were demultiplexed to FASTQ files using bcl2fastq2 (Illumina, v2.20). Sample quality was initially assessed using the BioSkryb BJ-DNA-QC (v2.0.5) pipeline implemented on the BaseJumper platform. Input libraries were downsampled to 40 million (40M) reads, after which the BJ-DNA-QC workflow performed an additional downsampling to 1 million read pairs.

Downsampling to 2 (2M) and 40 million read pairs was performed using seqtk (Li 2013 July 8) (v1.4). The downsampled reads as well as the full datasets for the MDA/REPLI-g samples were processed using Nextflow (Tommaso et al. 2017) (v24.4.1) through the nf-core/sarek (Garcia et al. 2020; Hanssen et al. 2024) workflow (v3.4.2). Read processing consisted of trimming with fastp (Chen et al. 2018; Chen 2023) (v0.23.4), mapping with BWA-MEM (Li and Durbin 2009) (v0.7.17.post1188) to reference genome GRCh38 (GCA_000001405.15_GRCh38_no_alt_analysis_set) and using GATK/picard (The picard toolkit development team 2019) (v4.5.0.0) for duplicate marking. MultiQC (Ewels et al. 2016) (v1.21) was used to aggregate statistics across samples.

Coverage depth and evenness analysis were performed using mosdepth (Pedersen and Quinlan 2017) (v0.3.8) using only reads with a minimum MAPQ of 20 with parameters “--no-per-base --fast-mode --mapq 20”. For the evenness analysis, mosdepth was run with a bin size of 100,000 bp with option “-b 100000”. To generate the Lorenz curve, bins were sorted by coverage in ascending order. The normalized cumulative coverage for each bin was then plotted against the normalized bin rank to generate the curve. The median absolute pairwise deviation (MAPD) was calculated as the median of the absolute coverage differences between pairs of neighboring bins. Only chromosomes 1 to X were considered for the evenness analysis.

Coverage breadth was evaluated using preseq (Daley and Smith 2014) (v3.1.2) with the “gc_extrap” command. For this purpose, properly paired reads were extracted while excluding supplementary reads using samtools (v1.19.2). Reads were converted from BAM to BED format using bedtools (Quinlan and Hall 2010) (v2.30.0) “bamtobed” command for input into preseq. To get the fraction of the genome covered, picard tools (v3.0.0) subcommand “NonNFastaSize” was used to calculate the genome size.

Samtools (Danecek et al. 2021) view (v1.19.2) was used to count chimeric reads, excluding unmapped and supplementary reads. To only keep confident mappings, reads with a MAPQ below 60 were excluded. Chimeric reads were defined as read pairs with mates on separate chromosomes or separated by more than 100,000 bp.

GC bias was investigated with picard tools (v3.0.0) using the “CollectGcBiasMetrics” command with default arguments. Briefly, the genome was broken into 100 bp windows and for each window the GC-rate and coverage was collected. For the plot, only GC-rates covered by at least 1,000,000 windows were included.

### Genotyping array data analysis

SNP coverage information was extracted using the Illumina Dragen Array software (v1.2.0). Briefly, genotypes were re-called from the IDAT files using the “dragena genotype call” command. These were then converted into bedgraph files containing LogR-ratio (LRR) and B-allele frequency (BAF) data for each SNP position using the “dragena genotype gtc-to-bedgraph” command. To get the average LRR per 100 kb, BEDtools (v2.30.0) was used. To calculate the corresponding ratios from the sequencing data, the read coverage was divided by the median coverage and then log2 transformed. For the genotype and SNV benchmarking analysis, the genotype call files (GTF) from Dragen Array were converted to VCF format using the “dragena genotype gtc-to-vcf” command.

For the benchmarking of the array SNV calls, we used a truth-set generated from PacBio HiFi sequencing using the extracted DNA from the same MM.1S cell line (Höjer et al. 2026). The PacBio reads mapped to GRCh38 using pbmm2 (v1.10.0, https://github.com/PacificBiosciences/pbmm2) are available on the European Nucleotide Archive ENA (PRJEB95775). Variants were called using DeepVariant (v1.5, “PACBIO” model), only keeping calls that passed default DeepVariant filters. Benchmarking was performed using hap.py (v0.3.15) (Krusche et al. 2019) with vcfeval (v3.12.1) and the analysis limited only to genome positions where array genotypes were called, and only substitutions were included in the final results.

### Copy number variant calling and benchmarking

Copy number variant (CNV) calling using Illumina single-cell and bulk sequencing data was performed using both CopyKit (Minussi et al. 2022) and Ginkgo (Garvin et al. 2015). CopyKit (v0.1.2, https://github.com/navinlabcode/copykit), which is configured for the reference genome GRCh38, was run with bin sizes of 110kb, 220kb (default), 500kb, and 1Mb. Copy numbers were inferred using the “fixed” method with a ploidy set to 2. Copy number segments were extracted and converted to SEG files using custom scripts for visualization using IGV (Robinson et al. 2011) and benchmarking. Ginkgo CNV (v0.0.2, https://github.com/robertaboukhalil/ginkgo) calling was run through the BaseJumper Bioinformatics Platform (BioSkryb), pipeline BJ-CNV (v1.2.4), with bin size 1Mb (default), and ploidy set to 2 (default).

For the CNV benchmarks, we again used the PacBio HiFi sequencing data. CNVs were called using HiFiCNV (v0.1.7, https://github.com/PacificBiosciences/HiFiCNV) excluding regions defined in https://github.com/PacificBiosciences/HiFiCNV/raw/refs/heads/main/data/excluded_regions/cnv.excluded_regions.common_50.hg38.bed.gz with option “--exclude cnv.excluded_regions.common_50.hg38.bed” following the developers’ recommendations.

SNP array CNVs were called with PennCNV (Wang et al. 2007) (commit 23576a3). Briefly, for each SNP signal values in the form of BAF and LRR values were extracted from the Dragen Array VCFs using bcftools (Danecek et al. 2021) (v1.20). GC correction values were generated from looking at the GC-rate in 1 Mb windows surrounding each SNP using BEDtools (v2.30.0). For the HMM model we used the configs in the “hhall.hmm” file, supplied with the PennCNV codebase. Using these files, raw CNVs were called using the PennCNV “detect_cnv.pl” script, including only CNVs covering at least 10 markers and a minimum length of 100kbp with options “--minsnp 10” and “--minlength 100000”. The PennCNV “clean_cnv.pl” script with the subcommand “combineseg” was then used to merge broken-up calls using option “--fraction 0.2”: This command merges CNVs with the same copy state if they are split by a gap of maximum 20% of their combined length.

CNV call performance against PacBio HiFiCNV calls were assessed by computing the basepair-level copy number F1 score. Briefly, for each base the assigned copy number was compared to the corresponding PacBio HiFiCNV copy number for agreement. Bases with the same copy number were tallied as true positives (TP). Disagreeing bases were tallied as (1) false negatives (FN) if HiFiCNV called neutral copy state (CN=2) or (2) false positive if HiFiCNV called a gain/loss (CN≠2). The F1 score was then calculated as follows:

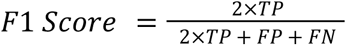

The CNV benchmarking was performed using in-house Python (v3.12.7) script using the pyranges (Stovner and Sætrom 2019) and pandas (McKinney 2010; The pandas development team 2020) (v2.2.3) packages.

### Data visualization

Data visualizations were generated using Python (v3.12.7) with extensive use of packages jupyter (Kluyver et al. 2016) (v5.7.2), scipy (Virtanen et al. 2020) (v1.14.1), seaborn (Waskom 2021) (v0.13.2), statsmodels (Skipper and Josef 2010) (v0.14.4), matplotlib (Hunter 2007) (v3.9.2), numpy (Harris et al. 2020) (v2.1.3), pandas (McKinney 2010; The pandas development team 2020) (v2.2.3). Figures and illustrations were composed and adjusted for visual clarity using Affinity Designer (v1.10.8 and v2.6.0).

### LLM statement

Gemini (3.5 Flash and 3.6 Flash, Google) and ChatGPT (GPT-5.6 Luna, OpenAI) LLMs were used to improve syntax and the legibility of the text as well as drafting some paragraphs/sentences. All text suggestions from the LLMs were reviewed to ensure the accuracy of the written text.

## RESULTS

### PTA/ResolveDNA outperforms MDA/REPLI-g across most quality metrics

PTA/ResolveDNA and MDA/REPLI-g were used to prepare single-cell and bulk (10 cells) libraries using cells from the MM.1S cancer cell line (Figure 1A). Ten single-cell libraries and one bulk were selected for sequencing from each kit, for a total of 22 libraries sequenced. Library yields were similar to the values reported by the manufacturers (Figure 1B). Library sequencing yielded ∼50 million paired reads for the PTA/ResolveDNA samples and 100 million or more for MDA/REPLI-g samples (Table 1).

**Table 1:** Sample read statistics. Read and alignment statistics for single-cell and bulk samples.

| Sample | Read pairs (M) | Mapped* | Duplicates* | Mean insert size (bp)* |
| --- | --- | --- | --- | --- |
| MDA/REPLI-g (Cell#01) | 238 | 99.98% | 0.6% | 219.6 |
| MDA/REPLI-g (Cell#02) | 185 | 99.99% | 0.7% | 220.6 |
| MDA/REPLI-g (Cell#03) | 165 | 99.98% | 0.8% | 217.5 |
| MDA/REPLI-g (Cell#04) | 169 | 99.98% | 0.8% | 214.6 |
| MDA/REPLI-g (Cell#05) | 138 | 99.98% | 0.8% | 217.3 |
| MDA/REPLI-g (Cell#06) | 185 | 99.98% | 0.7% | 214.8 |
| MDA/REPLI-g (Cell#07) | 234 | 99.99% | 0.6% | 215.7 |
| MDA/REPLI-g (Cell#08) | 212 | 99.99% | 0.6% | 218.6 |
| MDA/REPLI-g (Cell#09) | 175 | 99.98% | 0.6% | 220.7 |
| MDA/REPLI-g (Cell#10) | 110 | 99.98% | 1.1% | 215.9 |
| MDA/REPLI-g (Bulk) | 98 | 99.99% | 1.2% | 218.4 |
| PTA/ResolveDNA (Cell#01) | 55 | 99.98% | 1.1% | 208.4 |
| PTA/ResolveDNA (Cell#02) | 47 | 99.97% | 1.2% | 205.4 |
| PTA/ResolveDNA (Cell#03) | 57 | 99.97% | 1.1% | 202.6 |
| PTA/ResolveDNA (Cell#04) | 58 | 99.97% | 1.2% | 215.8 |
| PTA/ResolveDNA (Cell#05) | 64 | 99.97% | 0.9% | 203.5 |
| PTA/ResolveDNA (Cell#06) | 64 | 99.98% | 1.0% | 204.2 |
| PTA/ResolveDNA (Cell#07) | 63 | 99.97% | 1.0% | 199.7 |
| PTA/ResolveDNA (Cell#08) | 47 | 99.96% | 1.3% | 212.0 |
| PTA/ResolveDNA (Cell#09) | 56 | 99.97% | 1.0% | 204.5 |
| PTA/ResolveDNA (Cell#10) | 60 | 99.98% | 1.1% | 204.7 |
| PTA/ResolveDNA (Bulk) | 62 | 99.98% | 1.1% | 203.4 |
| *Based on samples downsampled to 40 million read pairs |  |  |  |  |

Initial library quality assessment using the BioSkryb BJ-DNA-QC pipeline shows that most (8/10) of the PTA/ResolveDNA samples fulfilled the validation metrics as defined in McDaniel et al. (McDaniel et al. 2025). For the MDA/REPLI-g, only the bulk library fulfilled the same criteria (Supplementary Table 1).

To further assess the sequencing libraries generated by both kits, samples were downsampled to 40M reads and aligned to GRCh38. All libraries showed high mapping rates (>99.9%) and low duplicate rates (<2%) (Table 1). The mean insert size was slightly above 200 bp, with MDA/REPLI-g libraries being marginally longer on average (Table 1).

The MDA/REPLI-g single cells displayed a wider range of coverage depth relative to the bulk and PTA/ResolveDNA samples and left a larger proportion of the genome uncovered (Figure 2A). For single cells, the Qiagen kit left on average 42% of the genome uncovered, compared to 25% for PTA/ResolveDNA. For reference, in the bulk samples ∼21% of the genome was uncovered for both kits. To further evaluate the coverage breadth of the samples, we extrapolated the expected coverage with up to 200 million reads (Figure 2B). Most PTA/ResolveDNA samples approach 100% coverage, while the MDA/REPLI-g samples lack coverage in around 15% of the genome.

**Figure 2.**
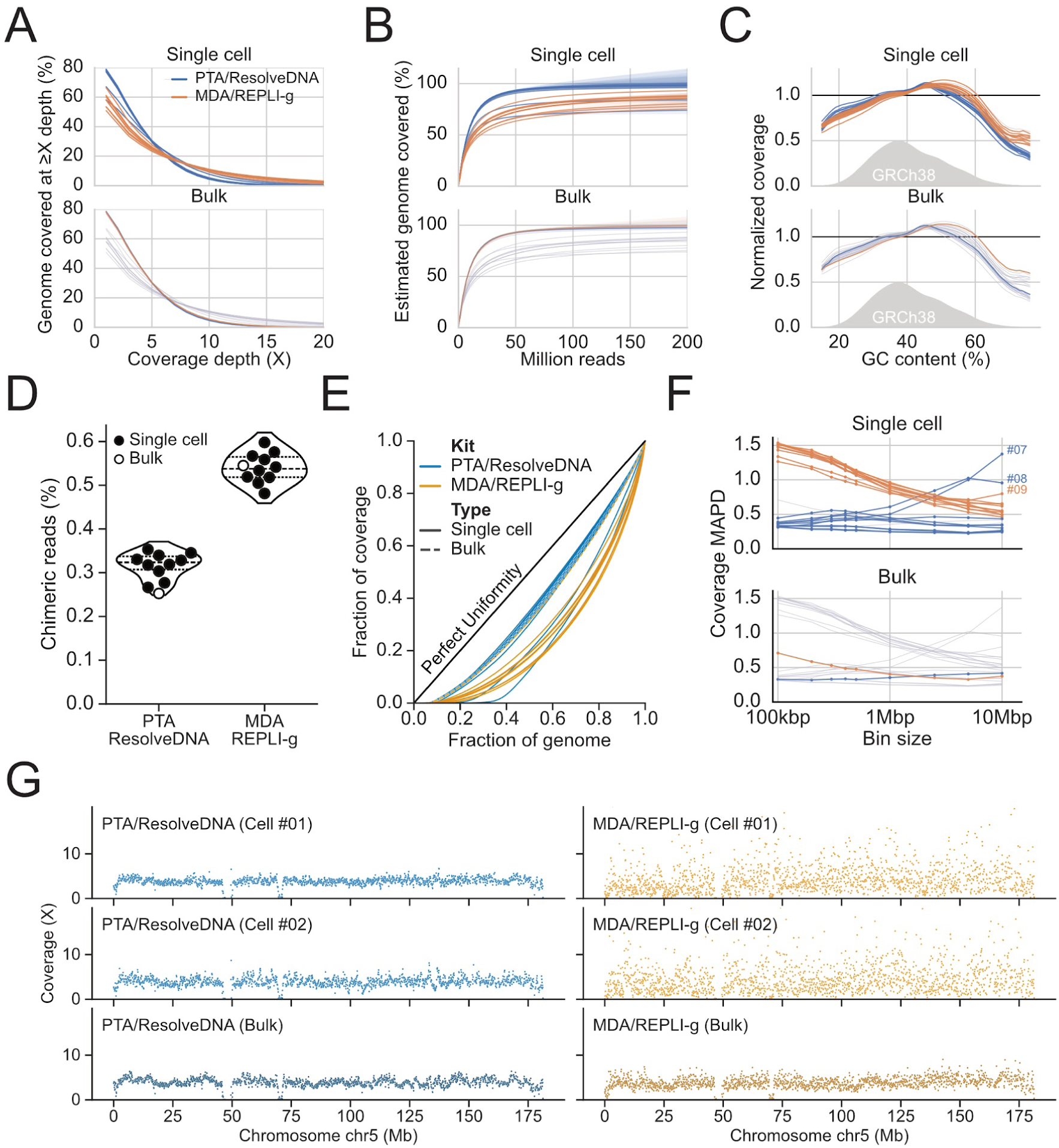
scWGS library characteristics for PTA/ResolveDNA and MDA/REPLI-g. All samples were subsampled to 40M reads. (**A**) Genome coverage distribution. (**B**) Percentage of genome covered as estimated from 40M reads using preseq gc_extrap. (**C**) GC-bias as the normalized coverage for regions of different GC-rate. The horizontal line at 1.0 represents unbiased coverage. (**D**) Percentage of mapped reads classified as being of chimeric origin. (**E**) Lorenz curve showing coverage evenness across 100 kb bins. (**F**) Median absolute pairwise deviation (MAPD) across bins from 100kb to 10 megabasepairs (Mb). (**G**) Examples showing coverage evenness across chromosome 5 in 100 kb bins.

DNA amplification is known to be affected by GC content. We therefore evaluated GC bias. While both kits showed similar low bias for most of the GC range found in the human genome, the MDA/REPLI-g displayed improved coverage in high GC content regions compared to PTA/ResolveDNA (Figure 2C).

MDA-based WGA is known to be associated with hyperbranching of DNA which is believed to increase issues such as the formation of chimeric DNA molecules (Lasken and Stockwell 2007). To evaluate this, we assessed chimeric formation by looking at split read pairs. Pairs with mates mapping with high confidence (MAPQ≥60) on separate chromosomes or spaced by more than 100,000 bp were classified as chimeric. For the single cells, 0.54% of the MDA/REPLI-g reads were on average classified as chimeric, compared to only 0.32% for PTA/ResolveDNA (Figure 2D). This result is in line with PTA not producing hyperbranched copies of the template DNA.

Uneven genome coverage leads to uncovered regions, hindering accurate copy number analysis. For this purpose, coverage evenness was evaluated using multiple metrics across the genome. Looking at the Lorenz curve, where a perfectly uniform coverage would follow the diagonal, PTA/ResolveDNA samples produce more uniform coverage, with only the MDA/REPLI-g bulk showing a similar distribution (Figure 2E). The Lorenz curve is affected by the underlying genome structure, where large duplications or deletions impact the perceived evenness. For this purpose, we also calculated the median absolute pairwise deviation (MAPD) which measures the coverage difference of neighboring bins. This produces a measure of local bin-to-bin variability with limited impact from genome differences. Across all bin sizes, the MAPD was overall significantly lower in the PTA/ResolveDNA (Figure 2F), indicating more local coverage evenness. Altogether, coverage evenness was better in the PTA/ResolveDNA samples. This could also be confirmed by visually inspecting the coverage across chromosome 5 (Figure 2G).

For two cells (#07 and #08) processed with the PTA/ResolveDNA, an elevated MAPD at larger bin sizes was noted (Figure 2F). These same cells did not pass the BJ-DNA-QC validation metrics (Supplementary Table 1) and also deviated from the other PTA/ResolveDNA samples in many other metrics (Figure 2A, 2B and 2E). WGA DNA yield for these samples did not deviate from the other scWGA PTA/ResolveDNA reactions, whose concentrations were all in the range for an evenly amplified genome (>15ng/μL in 40 μL of final reaction (Derks et al. 2025)). Further inspection revealed features consistent with ongoing replication, suggestive of S-phase cells (Supplementary Figure 1A). Interestingly, the two cells show very low pairwise correlation (Supplementary Figure 1B,C), which is in line with earlier findings (Chen et al. 2017) indicating stochastic replication firing. For completeness, these cells were included in the downstream analyses.

### PTA/ResolveDNA CNV calls are highly concordant to PacBio HiFi calls

To assess the performance of the tested kits in detecting CNVs, we called CNVs using Ginkgo and CopyKit across varying bin sizes and read depths. The results were benchmarked against HifiCNV-called CNVs identified from PacBio HiFi sequencing of the same cell line (Figure 3A). Coverage-based CNV callers such as Ginkgo and CopyKit output unprecise CNV boundaries and split CNVs which complicates CNV-to-CNV comparison. To get an estimate of overall CNV call performance, we computed the basepair copy number F1 score for each kit, caller and bin size compared to PacBio (see Methods for details). PTA/ResolveDNA generally had a high F1 score for almost all the single cells across callers and bin sizes even at low (2M) read depth (Figure 3B). In contrast, MDA/REPLI-g libraries exhibited only marginal improvements in CNV concordance at full sequencing depth (110–238 M reads; Figure 3B). Finally, the mean copy number F1 score for the PTA/ResolveDNA and MDA/REPLI-g bulk samples was 0.95 and 0.97, respectively.

**Figure 3:**
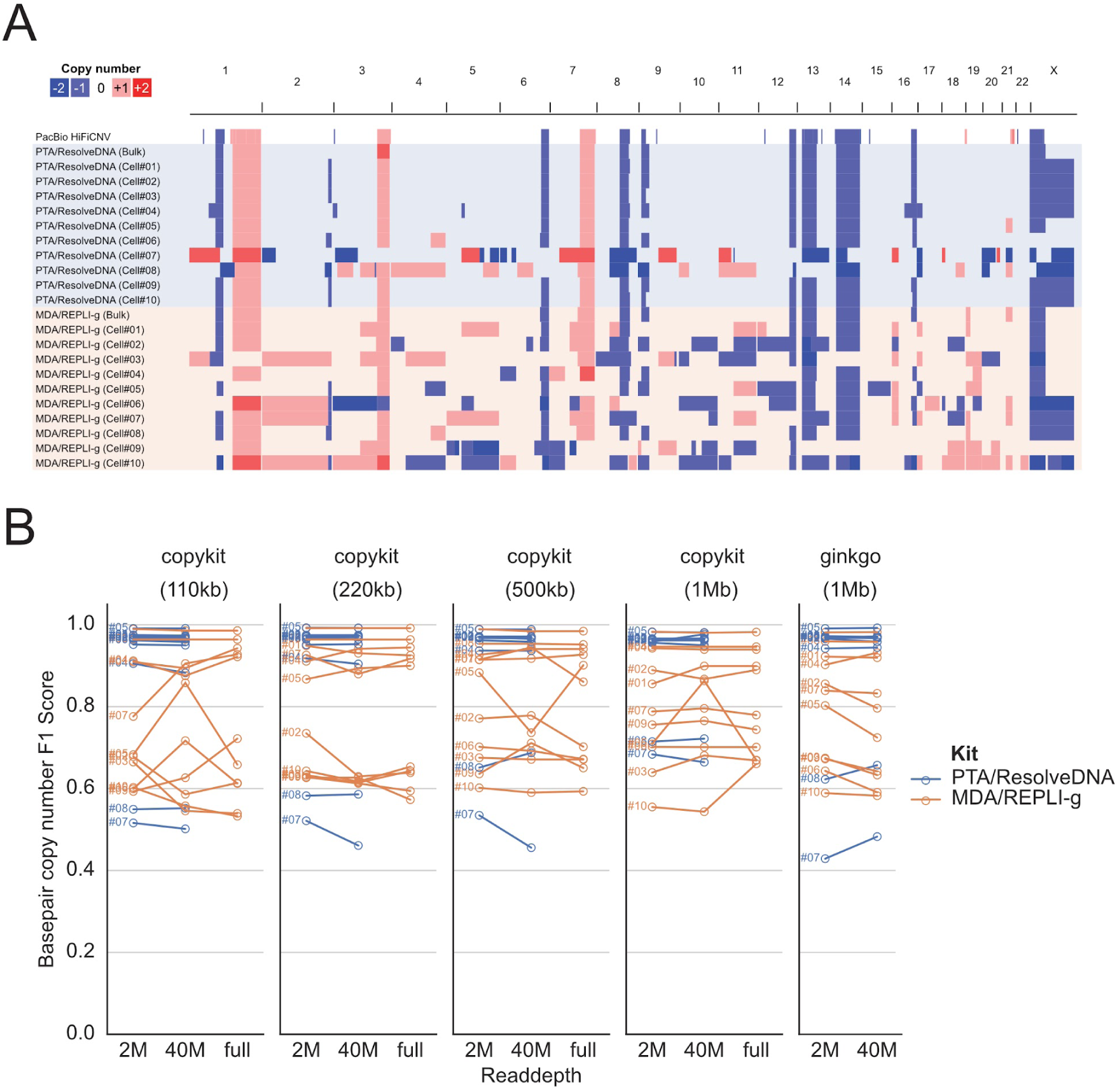
Single-cell CNV calling agreement. CNVs were called using CopyKit and Ginkgo at various read depths and with different resolutions. Calls were benchmarked against CNV calls from PacBio HiFi sequencing generated using the same cell line. (**A**) Genome-wide overview of copykit (1Mb bin size) CNVs using 40M reads. Image generated using IGV genome browser. (**B**) Basepair copy number F1 score as compared to bulk PacBio HiFiCNV calls. Read depth “full” refers to full read depth for the MDA/REPLI-g samples, which is in the range of 98-238 million reads (see Table 1 for actual numbers).

Given that the PTA/ResolveDNA kit demonstrated higher overall accuracy in detecting CNVs at the single-cell level, we next assessed its sensitivity for identifying smaller events. MDA-WGS has been suggested to be unsuitable for calling CNVs below 1Mb due to excessive coverage unevenness (Ning et al. 2015). To investigate this, we examined three representative CNVs (1–3 Mb) validated by PacBio HiFi sequencing. Single-cell CNVs were called with CopyKit using the smallest bin size (110 kb). All three CNVs were consistently detected in the majority of the PTA/ResolveDNA single-cell samples, even at the lowest read depth, whereas they were largely undetectable in the MDA/REPLI-g samples (Figure 4A). The pronounced coverage variability surrounding these loci in the MDA/REPLI-g data further illustrates the limitations of MDA for detecting small CNVs (Figure 4B). In addition, a 358 kb deletion on chromosome 3 was identified in the PacBio data (Figure 4A). Although this smaller CNV was not detected by the single-cell CNV callers evaluated, the coverage profiles (Figure 4B) indicate that it may be detectable in the PTA/ResolveDNA data with further parameter optimization.

**Figure 4:**
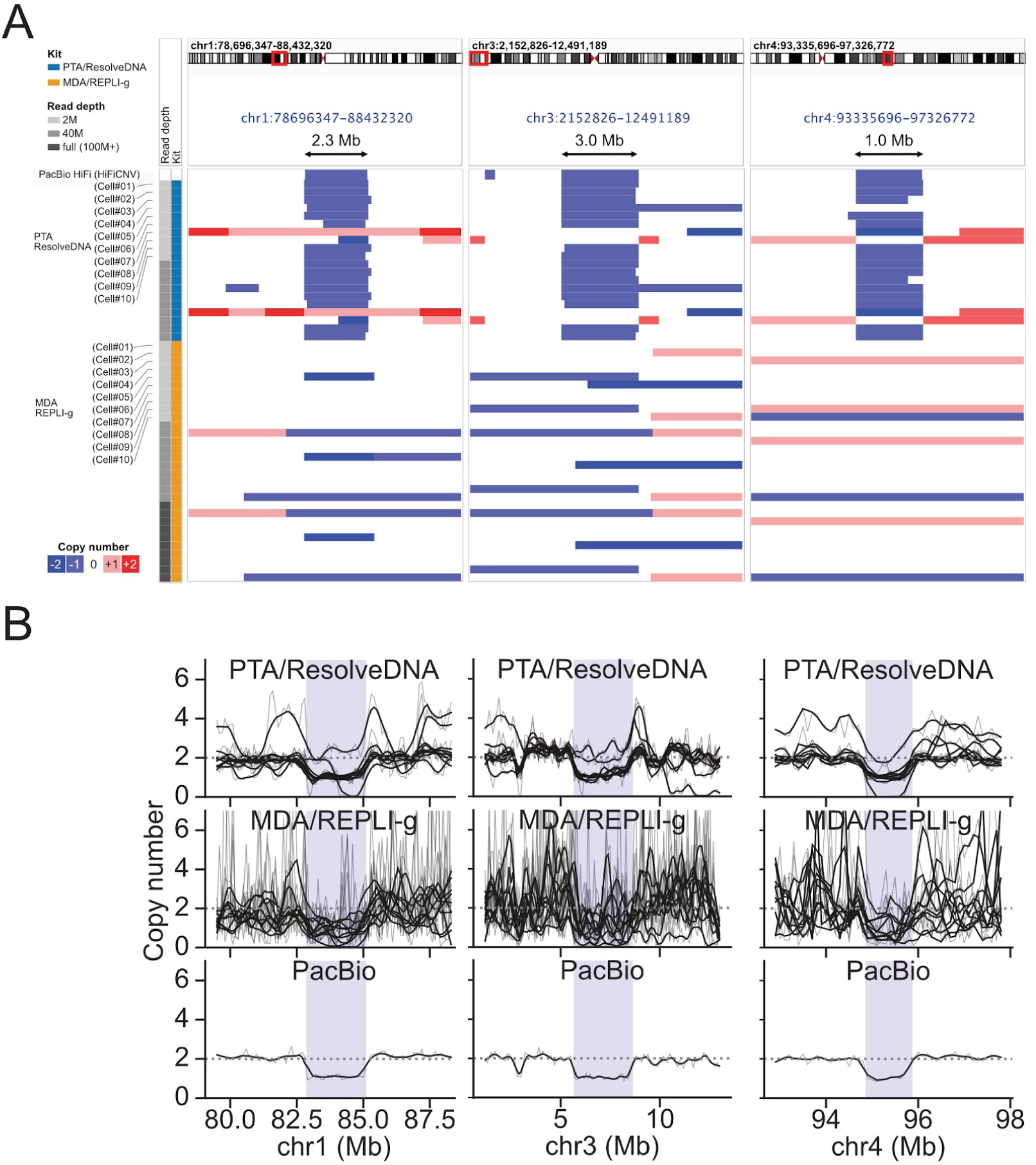
Small CNV calling sensitivity using three example CNVs (1-3Mb). (**A**) Single-cell CNV calls compared to PacBio calls. Single-cell CNVs were called using CopyKit with a bin size of 110kb. Image generated using IGV genome browser. (**B**) Normalized coverage from 40M reads surrounding the same CNVs as in A, compared to PacBio as highlighted in blue. Coverage was measured in 100 kb bins and normalized to half the median overall coverage to get the bin copy number. Raw bin copy number shown in gray and LOWESS smoothed values shown in black.

### Genotyping arrays as an option for single-cell WGA analysis

Next, we evaluated genome-wide SNP genotyping arrays (SNP arrays) for analysis of scWGA samples. Particularly for quality control, SNP arrays could be a cost-effective alternative to library preparation and low-pass sequencing. In addition, SNP arrays provide allelic level information that would otherwise require deep (30X) sequencing at considerably higher cost. Using the WGA DNA from the same two bulk and the 20 single-cell samples used for scWGS, we performed genotyping using the Illumina Infinium Global Screening Array (v3.0), which targets 650,181 variants across the human genome.

We first evaluated the calling rate for the variants included in the array (Figure 5A). The bulk and PTA/ResolveDNA single-cell samples have a high (>94%) call rate, except the two S-phase cells (#7 and #8) identified earlier. The MDA/REPLI-g single-cell samples had a lower mean call rate of 81%. This is in line with the coverage dropout observed in the sequencing-based analysis.

**Figure 5:**
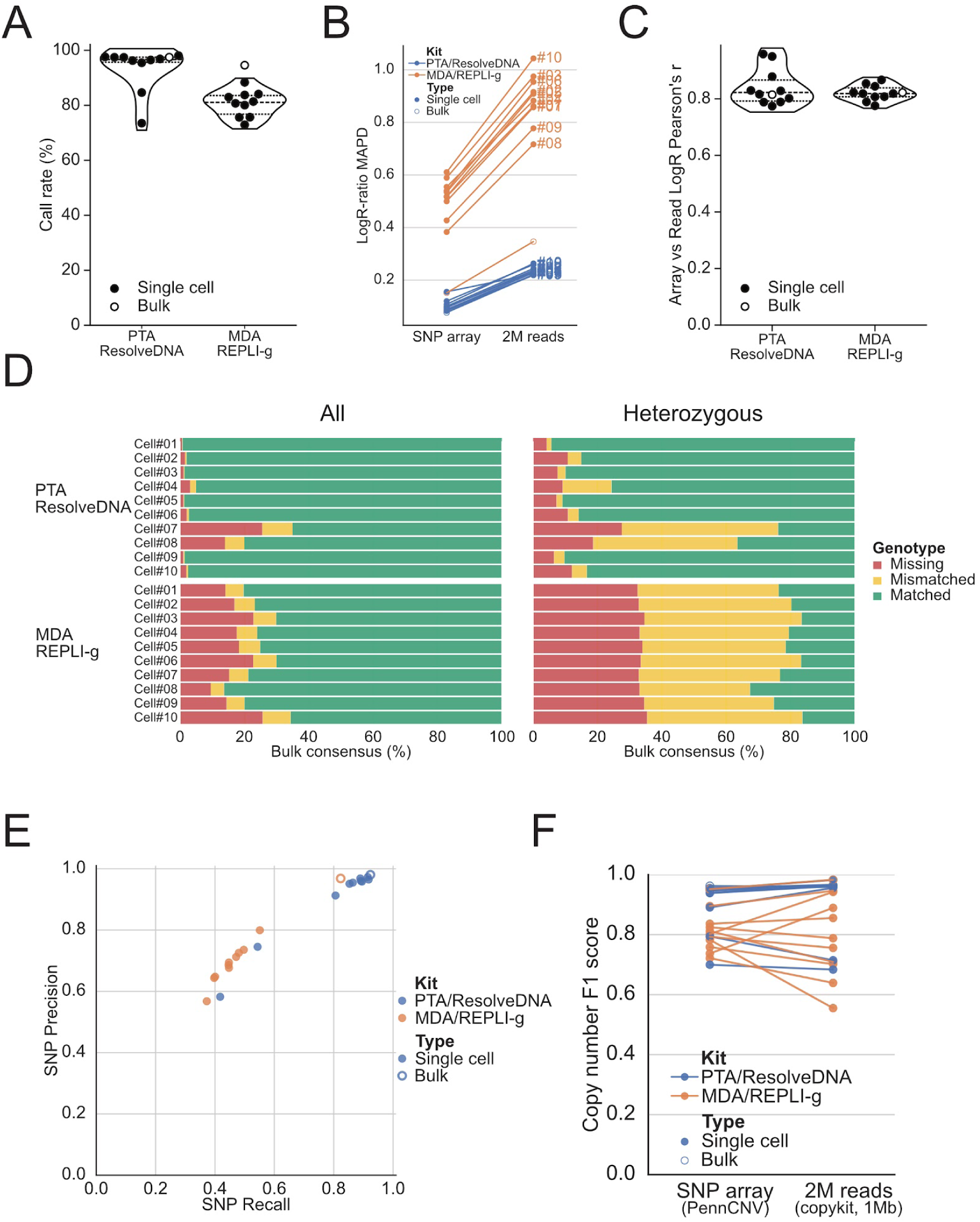
Genotyping SNP arrays for single-cell WGA DNA analysis. (**A**) SNP array call rate per kit for the single-cell and bulk samples. (**B**) Coverage evenness for SNP arrays versus low-pass sequencing (2M reads) as median absolute pairwise deviation (MAPD). The metrics were calculated from the LogR ratio (LRR) values in 100 kb bins. For the array data, the mean LRR was calculated in each bin. For the sequencing data, LRR was calculated using the log2 transform of the median normalized bin coverage. (**C**) Correlation between SNP array intensities and sequencing coverage LRR (**D**) Single-cell genotype concordance for bulk consensus genotypes using all genotypes (right) or only heterozygous genotypes (left). Markers without genotypes were classified as missing. (**E**) Precision and recall for single-cell array SNPs benchmarked to PacBio HiFi DeepVariant calls. (**F**) Copy number agreement compared to PacBio HiFiCNV truth-set. SNP array CNVs were called with PennCNV, while for the 2M reads CopyKit calls with 1 Mb bin size were used.

Next, we utilized the reported signal intensities from the SNP arrays to investigate WGA coverage uniformity. For this purpose, intensity LogR ratios (LRR) for each SNP position were computed and averaged across 100kb bins. For comparison, sequencing data downsampled to 2M reads was used to compute log-ratios across the same 100kb bins. To evaluate the coverage evenness, we calculated MAPD as earlier, which shows a similar trend with both the array intensities and low-pass sequencing data (Figure 5B). The array-based intensities correlated well with read coverage (mean Pearson’s correlation = 0.83; Figure 5C; Supplementary Figure 2).

We also used the array-based genotype information to assess the scWGA quality. Comparing the PTA/ResolveDNA and MDA/REPLI-g bulk, 94.3% of genotypes matched. Using these as a consensus set, we investigated the genotype concordance with the single cell samples. Looking at all genotype calls (n=602,083) the MDA/REPLI-g cells and PTA/ResolveDNA cells #07 and #08 were missing large fractions of calls (Figure 5D, left; Supplementary Table 2), which is in line with our above reported findings. Focusing on only heterozygous calls (n=64,038), ∼34% (21,539 on average) of calls were missing from the MDA/REPLI-g single-cell samples, with a ∼45% (28,636 on average) of calls having a mismatched genotype that indicates allelic dropout (Figure 5D, right; Supplementary Table 3).

Since MDA is commonly used for SNP calling we evaluated the genotype accuracy by comparing to PacBio DeepVariant called SNPs. The SNP genotype accuracy was again higher in the PTA/ResolveDNA data (Figure 5E), with the single cells displaying a median F1 score of 0.92. MDA/REPLI-g had noticeably lower recall in the single cells with a median recall of 0.45 compared to 0.88 for PTA/ResolveDNA.

Finally, we assessed the utility of SNP arrays in single-cell CNV calling. CNVs were called using PennCNV and benchmarked against the bulk PacBio HiFi calls. The sequencing-based CopyKit CNV calls with a 1Mb bin size using 2M reads were included for reference. While some larger variation in F1 score was observed for the MDA/REPLI-g single-cell samples, F1 score was generally high for the PTA/ResolveDNA PennCNV calls (median single-cell F1 score 0.94) with a level similar to the 2M reads (median single-cell F1 score 0.96) (Figure 5F, Supplementary Table 4).

## DISCUSSION

Single-cell whole genome analysis is a powerful technique for dissecting genetic heterogeneity at the cellular level. However, the necessity for amplification prior to analysis to overcome the limited DNA input from single cells introduces challenges, including amplification bias, uneven genome coverage, and artifacts that can compromise downstream analyses such as CNV and SNP detection.

In this study, we conducted a systematic comparison between the MDA/REPLI-g kit from Qiagen and the PTA/ResolveDNA kit from BioSkryb. WGA products were evaluated using both sequencing and genotype arrays. PTA/ResolveDNA outperformed MDA/REPLI-g in most metrics assessed, except for coverage in high GC regions. Genome coverage breadth and uniformity were markedly better in the PTA/ResolveDNA datasets. These findings are consistent with previous studies demonstrating the improved performance of PTA over conventional MDA-based approaches (Gonzalez-Pena et al. 2021; Kalef-Ezra et al. 2024). Nevertheless, it is important to note that MDA/REPLI-g has previously demonstrated high genome coverage breadth relative to several other non-PTA, scWGA kits (Estévez-Gómez et al. 2025).

The formation of chimeric reads, although relatively low in both methods, was elevated in the Qiagen libraries, consistent with the hyperbranching mechanism characteristic of MDA (Lasken and Stockwell 2007). This phenomenon, although subtle in frequency, can complicate structural variant analysis and should be considered with MDA-based protocols.

Two cells (#07 and #08) prepared with the PTA/ResolveDNA deviated from the rest across multiple metrics and showed indication of ongoing replication. Given the increased genome size of replicating cells, deviations in coverage breadth and evenness are expected. These findings suggest that PTA/ResolveDNA may also provide utility for studying replication dynamics, an application that has been challenging with MDA-based approaches (Aa et al. 2013).

The performance improvement of PTA/ResolveDNA was most evident in CNV detection, where PTA/ResolveDNA showed greater concordance with reference PacBio HiFiCNV calls, also capturing small CNVs in the 1–3 Mb range. These events were reliably detected in PTA/ResolveDNA samples even at low read depth or with genotyping arrays, whereas MDA/REPLI-g libraries largely failed to resolve these regions due to high local coverage variability. Earlier benchmarks have also observed a low CNV calling sensitivity for the REPLI-g kit (Deleye et al. 2017). In line with our results, MDA amplification bias impedes accurate CNV calling even when performed in nano- or picoliter droplets (dMDA) (Kalef-Ezra et al. 2024).

Genotyping SNP array-based analysis corroborated the sequencing findings. Call rates, coverage uniformity, and genotype concordance were consistently higher in PTA/ResolveDNA samples. The SNP arrays also allowed us to assess heterozygous genotype dropout rates, which would otherwise require deep sequencing. The dropout was indicative of allelic bias and poor amplification in the MDA/REPLI-g samples, reinforcing the limitations of MDA for comprehensive single-cell genotyping as reported elsewhere (Gonzalez-Pena et al. 2021). We observed that PTA/ResolveDNA CNV calling performance by SNP arrays was similar to low-pass sequencing, altogether highlighting SNP arrays as an attractive time/cost efficient option for single-cell genome analysis.

Several limitations in this study warrant consideration. First, our technological scope was restricted to PTA/ResolveDNA and MDA/REPLI-G, which both employ a similar WGA approach. We have therefore excluded alternative technologies, such as Ampli-1 and PicoPlex, which have performed well in other benchmarks (Kalef-Ezra et al. 2024; Estévez-Gómez et al. 2025). Second, our baseline assessment relied on a modest sample size of ten single cells per kit from a single multiple myeloma cell line (MM1S). Although MM.1S provides a highly dynamic genomic landscape ideal for evaluating copy number variant (CNV) detection, these findings may not perfectly generalize to normal diploid cells or other tumor types.

In this study, we used short read sequencing and SNP arrays approaches for the analyses of WGA DNA. Recently, long read sequencing has been introduced to enable structural variant detection at single-cell resolution, employing either droplet-based MDA (Hård et al. 2023) or Tn5-based WGA (Fan et al. 2021). While current PTA products yield inserts that are generally too short for optimal long-read sequencing, adapting this method for longer fragments could offer an even more comprehensive single-cell genome characterization. Indeed, long read PTA adaptation with the BioSkryb ResolveSeq kit is already showing promise for long-read scWGS (Swanson et al. 2025).

Our results reinforce the PTA/ResolveDNA kit advantages over the MDA/REPLI-g kit for single-cell WGA. These advantages include improved genome coverage, reduced amplification bias, enhanced CNV detection, and greater SNP genotyping fidelity. While both kits are capable of generating usable single-cell libraries, the performance differences are substantial, particularly for applications requiring CNV analysis and/or accurate genotyping. Furthermore, our results emphasize genotyping SNP arrays as a viable and practical tool alongside sequencing for quality control and CNV analysis of PTA single-cell WGA samples.

## ACKNOWLEDGMENTS

NGI is supported by the Swedish Research Council, SciLifeLab, the Knut and Alice Wallenberg Foundation, and ALF funding from Region Uppsala. Additional financial support was provided by the Swedish Blood Cancer Association, the Swedish Cancer Society, and Radiumhemmets forskningsfonder. We thank Joakim Lundeberg for his valuable input and discussions on the study design.

## DATA AND CODE AVAILABILITY

Illumina sequencing data for the scWGS kit samples is available on ENA under the study accession <u>PRJEB97702</u>. Genotyping array data is available on FigShare (DOI: https://doi.org/10.17044/scilifelab.30084184). PacBio HiFi BAM files for the MM.1S cell line are available on ENA under the study accession <u>PRJEB95775</u>. Analysis code (notebooks, workflow files, configs, scripts, etc) along with some data used to generate the figures are available on GitHub (<u>NationalGenomicsInfrastructure/NGI_scWGS_technote_2025</u>) with a persistent copy on Zenodo (DOI: https://doi.org/10.5281/zenodo.17296102).

## Notes

https://www.ebi.ac.uk/ena/browser/view/PRJEB97702

https://researchdata.se/en/catalogue/dataset/doi-10-17044-scilifelab-30084184

https://www.ebi.ac.uk/ena/browser/view/PRJEB95775

https://github.com/NationalGenomicsInfrastructure/NGI_scWGS_technote_2025

### Competing Interest Statement

The authors have declared no competing interest.

### Summary of Updates

We have reframed the scope and discussion so it is clear that PTA outperforming MDA is not a novel finding, Additionally we have: - Updated CNV analysis (F1 scores instead). - Including array SNV benchmarks against the PacBio data. - Referring to the kits instead of companies in the text/figures (PTA/ResolveDNA and MDA/REPLI-g).

