## Supplementary material for "Comparative analysis of whole genome amplification kits for single-cell genome analysis"

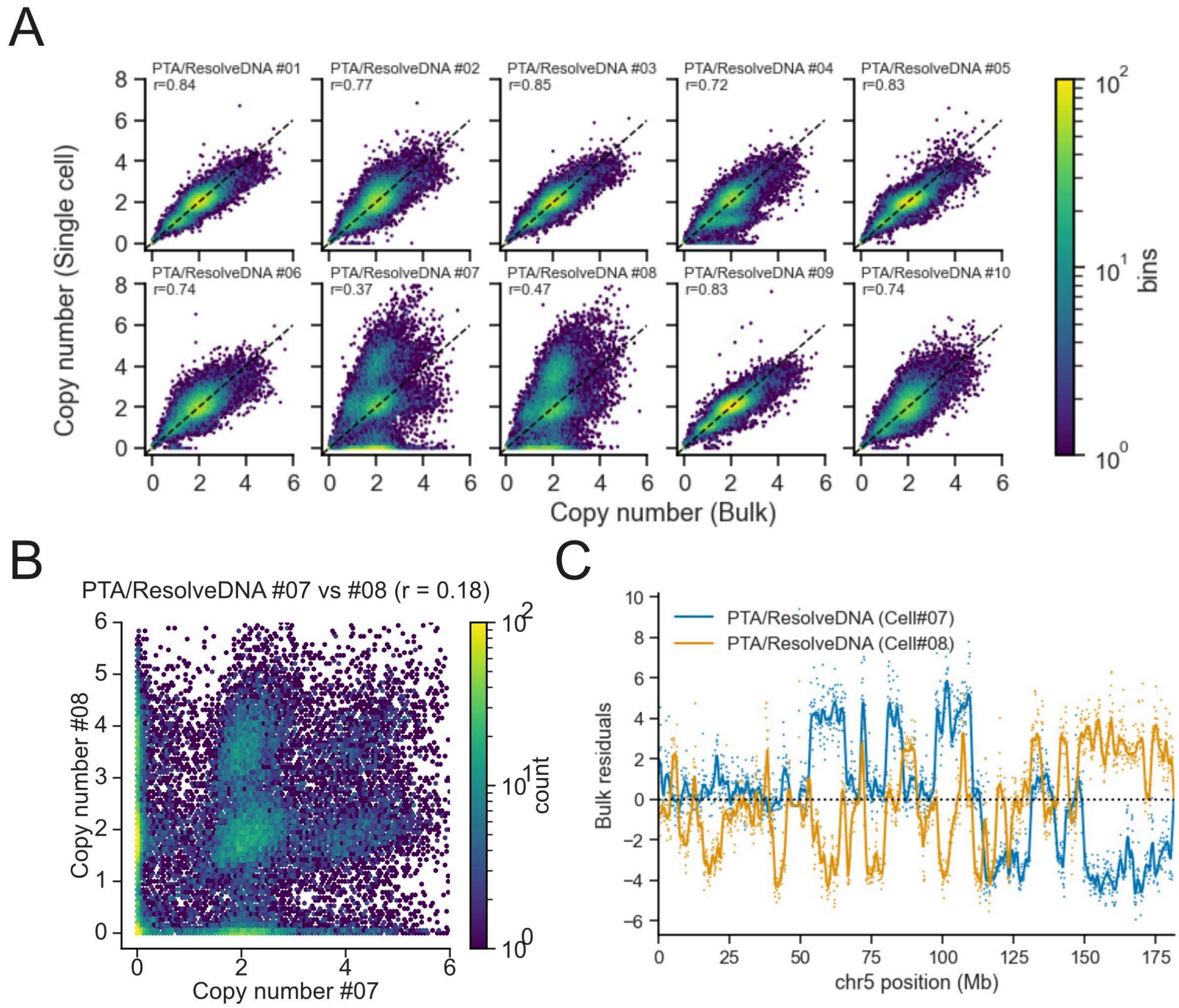


**Supplementary Figure 1: Evidence of S-phase characteristics in PTA/ResolveDNA cell #7 and #8.** (**A**) Bin copy number correlation comparing the single-cells to the bulk PTA/ResolveDNA samples. Copy numbers for 100kb bins are calculated as twice the median-normalized coverage. (**B**) Bin copy number correlation between cell #7 and #8. (**C**) Residuals after linear regression to the bulk along chromosome 5 for cell #7 and #8.

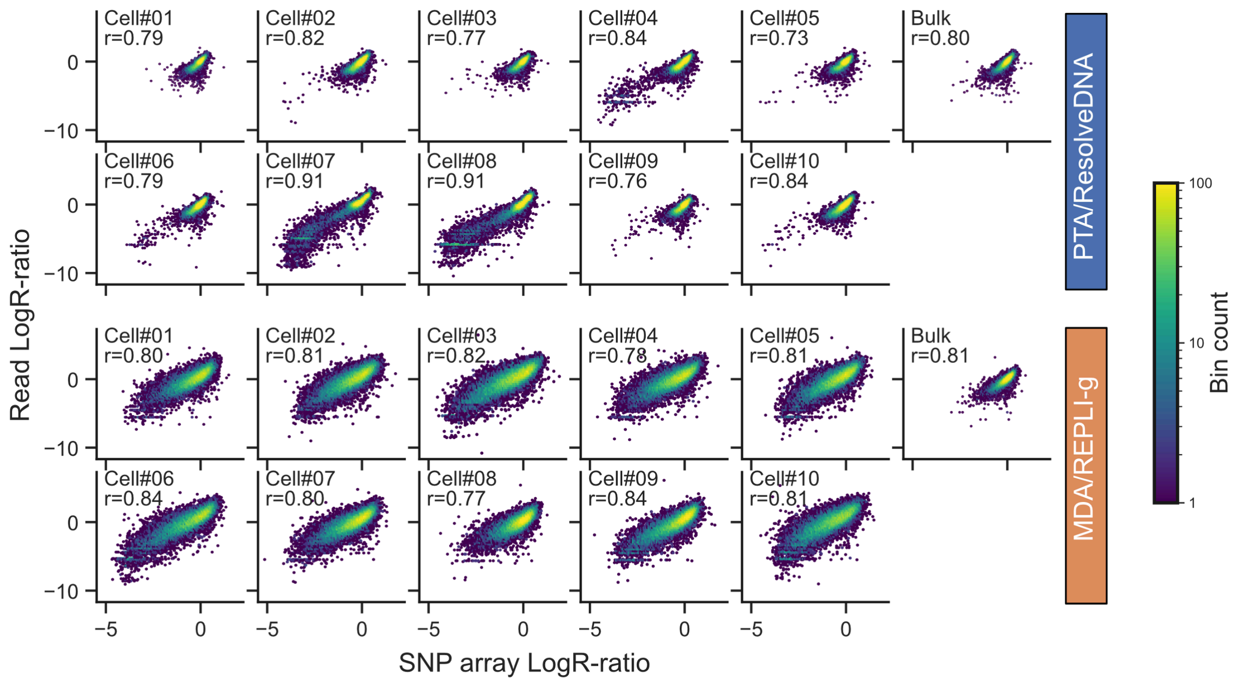


**Supplementary Figure 2:** Correlation between SNP array LogR-ratios and downsampled (2M) read coverage in 100kb bins. Read coverage converted to LogR-ratio by taking the log2 of the median normalized coverage. For each sample pair the Pearson correlation coefficient (r) is shown.

**Supplementary Table 1:** BJ-DNA-QC metrics. Threshold values were taken from McDaniel et al.(2025)

| Sample | PreSeq Count | % chrM | % Chimeras | Gini coefficient | Pass thresholds |
|---|---:|---:|---:|---:|---:|
| MDA/REPLI-g (Cell#01) | 2,419,053,312 | 0.06 | 4.8 | 0.107 | No |
| MDA/REPLI-g (Cell#02) | 2,258,415,116 | 0.01 | 5.0 | 0.108 | No |
| MDA/REPLI-g (Cell#03) | 2,240,058,661 | 0.04 | 5.0 | 0.111 | No |
| MDA/REPLI-g (Cell#04) | 2,183,339,638 | 0.02 | 5.2 | 0.109 | No |
| MDA/REPLI-g (Cell#05) | 2,374,487,162 | 0.06 | 5.0 | 0.109 | No |
| MDA/REPLI-g (Cell#06) | 1,998,460,660 | 0.01 | 4.9 | 0.110 | No |
| MDA/REPLI-g (Cell#07) | 2,385,796,247 | 0.04 | 4.8 | 0.106 | No |
| MDA/REPLI-g (Cell#08) | 2,846,911,943 | 0.04 | 4.5 | 0.103 | No |
| MDA/REPLI-g (Cell#09) | 2,406,294,721 | 0.02 | 5.0 | 0.106 | No |
| MDA/REPLI-g (Cell#10) | 1,929,595,118 | 0.01 | 5.2 | 0.114 | No |
| MDA/REPLI-g (Bulk) | 4,107,401,209 | 0.07 | 4.4 | 0.097 | Yes |
| PTA/ResolveDNA (Cell#01) | 3,902,855,314 | 0.31 | 6.4 | 0.098 | Yes |
| PTA/ResolveDNA (Cell#02) | 4,048,915,196 | 0.21 | 6.6 | 0.098 | Yes |
| PTA/ResolveDNA (Cell#03) | 4,091,147,708 | 0.29 | 6.4 | 0.098 | Yes |
| PTA/ResolveDNA (Cell#04) | 3,614,888,417 | 0.32 | 7.0 | 0.098 | Yes |
| PTA/ResolveDNA (Cell#05) | 4,319,861,697 | 0.31 | 6.6 | 0.098 | Yes |
| PTA/ResolveDNA (Cell#06) | 4,090,686,164 | 0.32 | 6.6 | 0.098 | Yes |
| PTA/ResolveDNA (Cell#07) | 2,571,765,384 | 0.25 | 6.9 | 0.104 | No |
| PTA/ResolveDNA (Cell#08) | 3,055,625,263 | 0.48 | 7.1 | 0.101 | No |
| PTA/ResolveDNA (Cell#09) | 4,168,932,266 | 0.27 | 6.7 | 0.098 | Yes |
| PTA/ResolveDNA (Cell#10) | 3,832,091,684 | 0.27 | 6.5 | 0.098 | Yes |
| PTA/ResolveDNA (Bulk) | 4,019,942,736 | 0.27 | 5.6 | 0.097 | Yes |
| Threshold value | >3,000,000,000 | <2.00 | <20.0 | <0.100 | |

**Supplementary Table 2**: Single-cell sample genotype concordance with mini-bulk consensus genotypes (n = 602,083) from Infinium GSAv3.0 array data.

| SampleName | Matched | % | Mismatched | % | Missing | % |
|---|---:|---:|---:|---:|---:|---:|
| PTA/ResolveDNA (Cell#01) | 597,384 | 99.2 | 952 | 0.2 | 3,747 | 0.6 |
| PTA/ResolveDNA (Cell#02) | 589,753 | 98.0 | 2,803 | 0.5 | 9,527 | 1.6 |
| PTA/ResolveDNA (Cell#03) | 593,772 | 98.6 | 1,673 | 0.3 | 6,638 | 1.1 |
| PTA/ResolveDNA (Cell#04) | 572,135 | 95.0 | 10,958 | 1.8 | 18,990 | 3.2 |
| PTA/ResolveDNA (Cell#05) | 594,204 | 98.7 | 1,453 | 0.2 | 6,426 | 1.1 |
| PTA/ResolveDNA (Cell#06) | 585,474 | 97.2 | 3,934 | 0.7 | 12,675 | 2.1 |
| PTA/ResolveDNA (Cell#07) | 391,675 | 65.1 | 56,615 | 9.4 | 153,793 | 25.5 |
| PTA/ResolveDNA (Cell#08) | 481,786 | 80.0 | 36,133 | 6.0 | 84,164 | 14.0 |
| PTA/ResolveDNA (Cell#09) | 593,605 | 98.6 | 2,246 | 0.4 | 6,232 | 1.0 |
| PTA/ResolveDNA (Cell#10) | 586,960 | 97.5 | 3,210 | 0.5 | 11,913 | 2.0 |
| MDA/REPLI-g (Cell#01) | 483,149 | 80.2 | 33,869 | 5.6 | 85,065 | 14.1 |
| MDA/REPLI-g (Cell#02) | 462,198 | 76.8 | 37,911 | 6.3 | 101,974 | 16.9 |
| MDA/REPLI-g (Cell#03) | 422,379 | 70.2 | 42,396 | 7.0 | 137,308 | 22.8 |
| MDA/REPLI-g (Cell#04) | 457,573 | 76.0 | 38,434 | 6.4 | 106,076 | 17.6 |
| MDA/REPLI-g (Cell#05) | 452,240 | 75.1 | 39,772 | 6.6 | 110,071 | 18.3 |
| MDA/REPLI-g (Cell#06) | 421,577 | 70.0 | 43,684 | 7.3 | 136,822 | 22.7 |
| MDA/REPLI-g (Cell#07) | 474,426 | 78.8 | 35,710 | 5.9 | 91,947 | 15.3 |
| MDA/REPLI-g (Cell#08) | 519,712 | 86.3 | 24,800 | 4.1 | 57,571 | 9.6 |
| MDA/REPLI-g (Cell#09) | 481,540 | 80.0 | 33,460 | 5.6 | 87,083 | 14.5 |
| MDA/REPLI-g (Cell#10) | 395,059 | 65.6 | 52,512 | 8.7 | 154,512 | 25.7 |

**Supplementary Table 3:** Single-cell sample genotype concordance with mini-bulk heterozygous consensus genotypes (n = 64,038) from Infinium GSAv3.0 array data.

| SampleName | Matched | % | Mismatched | % | Missing | % |
|---|---:|---:|---:|---:|---:|---:|
| PTA/ResolveDNA (Cell#01) | 60,409 | 94.3 | 914 | 1.4 | 2,715 | 4.2 |
| PTA/ResolveDNA (Cell#02) | 54,483 | 85.1 | 2,656 | 4.1 | 6,899 | 10.8 |
| PTA/ResolveDNA (Cell#03) | 57,578 | 89.9 | 1,573 | 2.5 | 4,887 | 7.6 |
| PTA/ResolveDNA (Cell#04) | 48,413 | 75.6 | 9,747 | 15.2 | 5,878 | 9.2 |
| PTA/ResolveDNA (Cell#05) | 58,196 | 90.9 | 1,171 | 1.8 | 4,671 | 7.3 |
| PTA/ResolveDNA (Cell#06) | 54,981 | 85.9 | 2,116 | 3.3 | 6,941 | 10.8 |
| PTA/ResolveDNA (Cell#07) | 15,305 | 23.9 | 31,073 | 48.5 | 17,660 | 27.6 |
| PTA/ResolveDNA (Cell#08) | 23,408 | 36.6 | 28,673 | 44.8 | 11,957 | 18.7 |
| PTA/ResolveDNA (Cell#09) | 57,781 | 90.2 | 2,085 | 3.3 | 4,172 | 6.5 |
| PTA/ResolveDNA (Cell#10) | 53,312 | 83.3 | 2,972 | 4.6 | 7,754 | 12.1 |
| MDA/REPLI-g (Cell#01) | 15,190 | 23.7 | 28,041 | 43.8 | 20,807 | 32.5 |
| MDA/REPLI-g (Cell#02) | 12,662 | 19.8 | 30,344 | 47.4 | 21,032 | 32.8 |
| MDA/REPLI-g (Cell#03) | 10,632 | 16.6 | 31,241 | 48.8 | 22,165 | 34.6 |
| MDA/REPLI-g (Cell#04) | 13,169 | 20.6 | 29,665 | 46.3 | 21,204 | 33.1 |
| MDA/REPLI-g (Cell#05) | 13,799 | 21.5 | 28,486 | 44.5 | 21,753 | 34.0 |
| MDA/REPLI-g (Cell#06) | 10,736 | 16.8 | 31,864 | 49.8 | 21,438 | 33.5 |
| MDA/REPLI-g (Cell#07) | 14,930 | 23.3 | 28,085 | 43.9 | 21,023 | 32.8 |
| MDA/REPLI-g (Cell#08) | 20,893 | 32.6 | 21,966 | 34.3 | 21,179 | 33.1 |
| MDA/REPLI-g (Cell#09) | 16,149 | 25.2 | 25,789 | 40.3 | 22,100 | 34.5 |
| MDA/REPLI-g (Cell#10) | 10,470 | 16.3 | 30,881 | 48.2 | 22,687 | 35.4 |

**Supplementary Table 4:** CNV F1 score compared to PacBio HiFiCNV calls for SNP array data with PennCNV caller and low pass (2M reads) sequencing with CopyKit caller (1Mb bins).

| Sample | SNP array (PennCNV) | 2M reads (CopyKit, 1Mb) |
|---|---:|---:|
| PTA/ResolveDNA (Bulk) | 0.95 | 0.96 |
| PTA/ResolveDNA (Cell#01) | 0.96 | 0.96 |
| PTA/ResolveDNA (Cell#02) | 0.96 | 0.95 |
| PTA/ResolveDNA (Cell#03) | 0.96 | 0.95 |
| PTA/ResolveDNA (Cell#04) | 0.96 | 0.89 |
| PTA/ResolveDNA (Cell#05) | 0.98 | 0.95 |
| PTA/ResolveDNA (Cell#06) | 0.96 | 0.94 |
| PTA/ResolveDNA (Cell#07) | 0.75 | 0.70 |
| PTA/ResolveDNA (Cell#08) | 0.76 | 0.79 |
| PTA/ResolveDNA (Cell#09) | 0.96 | 0.95 |
| PTA/ResolveDNA (Cell#10) | 0.96 | 0.94 |
| MDA/REPLI-g (Bulk) | 0.98 | 0.95 |
| MDA/REPLI-g (Cell#01) | 0.87 | 0.84 |
| MDA/REPLI-g (Cell#02) | 0.90 | 0.74 |
| MDA/REPLI-g (Cell#03) | 0.69 | 0.72 |
| MDA/REPLI-g (Cell#04) | 0.94 | 0.80 |
| MDA/REPLI-g (Cell#05) | 0.78 | 0.81 |
| MDA/REPLI-g (Cell#06) | 0.69 | 0.76 |
| MDA/REPLI-g (Cell#07) | 0.87 | 0.82 |
| MDA/REPLI-g (Cell#08) | 0.94 | 0.90 |
| MDA/REPLI-g (Cell#09) | 0.81 | 0.79 |
| MDA/REPLI-g (Cell#10) | 0.64 | 0.78 |
